# RNA Sequencing by Direct Tagmentation of RNA/DNA Hybrids

**DOI:** 10.1101/843474

**Authors:** Lin Di, Yusi Fu, Yue Sun, Jie Li, Lu Liu, Jiacheng Yao, Guanbo Wang, Yalei Wu, Kaiqin. Lao, Raymond W. Lee, Genhua Zheng, Jun Xu, Juntaek Oh, Dong Wang, X. Sunney Xie, Yanyi Huang, Jianbin Wang

## Abstract

Transcriptome profiling by RNA sequencing (RNA-seq) has been widely used to characterize cellular status but it relies on second strand cDNA synthesis to generate initial material for library preparation. Here we use bacterial transposase Tn5, which has been increasingly used in various high-throughput DNA analyses, to construct RNA-seq libraries without second strand synthesis. We show that Tn5 transposome can randomly bind RNA/DNA heteroduplexes and add sequencing adapters onto RNA directly after reverse transcription. This method, Sequencing HEteRo RNA-DNA-hYbrid (SHERRY), is versatile and scalable. SHERRY accepts a wide range of starting materials, from bulk RNA to single cells. SHERRY offers a greatly simplified protocol, and produces results with higher reproducibility and GC uniformity compared with prevailing RNA-seq methods.

**Significance Statement:** RNA sequencing is widely used to measure gene expression in biomedical research; therefore, improvements in the simplicity and accuracy of the technology are desirable. All existing RNA sequencing methods rely on the conversion of RNA into double-stranded DNA through reverse transcription followed by second strand synthesis. The latter step requires additional enzymes and purification, and introduces sequence-dependent bias. Here, we show that Tn5 transposase, which randomly binds and cuts double-stranded DNA, can directly fragment and prime the RNA/DNA heteroduplexes generated by reverse transcription. The primed fragments are then subject to PCR amplification. This provides a new approach for simple and accurate RNA characterization and quantification.

## Introduction

Transcriptome profiling through RNA sequencing (RNA-seq) has become routine in biomedical research since the popularization of next-generation sequencers and the dramatic fall in the cost of sequencing. RNA-seq has been widely used in addressing various biological questions, from exploring the pathogenesis of disease (1, 2) to constructing transcriptome maps for various species (3, 4). RNA-seq provides informative assessments of samples, especially when heterogeneity in a complex biological system (5, 6) or time-dependent dynamic processes are being investigated (7–9). A typical RNA-seq experiment requires a DNA library generated from mRNA transcripts. The commonly used protocols contain a few key steps, including RNA extraction, poly-A selection or ribosomal RNA depletion, reverse transcription, second strand cDNA synthesis, adapter ligation, and PCR amplification (10–12).

Although many experimental protocols, combining novel chemistry and processes, have recently been invented, RNA-seq is still a challenging technology to apply. On one hand, most of these protocols are designed for conventional bulk samples (11, 13–15), which typically contain millions of cells or more. However, many cutting-edge studies require transcriptome analyses of very small amounts of input RNA, for which most large-input protocols do not work. The main reason of this incompatibility is because the purification operations needed between the main experimental steps cause inevitable loss of the nucleic acid molecules.

On the other hand, many single-cell RNA-seq protocols have been invented in the past decade (16–19). However, for most of these protocols it is difficult to achieve both high throughput and high detectability. One type of single-cell RNA-seq approach, such as Smart-seq2 (17), is to introduce pre-amplification to address the low-input problem but such an approach is likely to introduce bias and to impair quantification accuracy. Another type of approach is to barcode each cell’s transcripts and hence bioinformatically assign identity to the sequencing data that is linked to each cell and each molecule (20–23). However, the detectability and reproducibility of such approaches are still not ideal (24). An easy and versatile RNA-seq method is needed that works with input from single cells to bulk RNA.

Bacterial transposase Tn5 (25) has been employed in next generation sequencing, taking advantage of the unique ‘tagmentation’ function of dimeric Tn5, which can cut double-stranded DNA (dsDNA) and ligate the resulting DNA ends to specific adaptors. Genetically engineered Tn5 is now widely used in sequencing library preparation for its rapid processivity and low sample input requirement (26, 27). For general library preparation, Tn5 directly reacts with naked dsDNA. This is followed by PCR amplification with sequencing adaptors. Such a simple one-step tagmentation reaction has greatly simplified the experimental process, shortening the workflow time and lowering costs. Tn5-tagmentation has also been used for the detection of chromatin accessibility, high-accuracy single-cell whole-genome sequencing, and chromatin interaction studies (28–32). For RNA-seq, the RNA transcripts have to undergo reverse transcription and second strand synthesis, before the Tn5 tagmentation of the resulting dsDNA (33, 34).

In this paper we present a novel RNA-seq method using Tn5 transposase to directly tagment RNA/DNA hybrids to form an amplifiable library. We experimentally show that, as an RNase H superfamily member (35), Tn5 binds to RNA/DNA heteroduplex similarly as to dsDNA and effectively fragments and then ligates the specific amplification and sequencing adaptor onto the hybrid. This method, named Sequencing HEteRo RNA-DNA-hYbrid (SHERRY), greatly improves the robustness of low-input RNA-seq with a simplified experimental procedure. We also show that SHERRY works with various amounts of input sample, from single cells to bulk RNA, with a dynamic range spanning six orders of magnitude. SHERRY shows superior cross-sample robustness and comparable detectability for both bulk RNA and single cells compared with other commonly used methods and provides a unique solution for small bulk samples that existing approaches struggle to handle. Furthermore, this easy-to-operate protocol is scalable and cost effective, holding promise for use in high-quality and high-throughput RNA-seq applications.

## Results

### New RNA-seq strategy using RNA/DNA hybrid tagmentation

Because of its nucleotidyl transfer activity, transposase Tn5 has been widely used in recently developed DNA sequencing technologies. Previous studies (36, 37) have identified a catalytic site within its DDE domain (**Fig. S1A)**. Indeed, when we mutated one of the key residues (D188E) (38) in pTXB1 Tn5, its fragmentation activity on dsDNA was notably impaired (**Fig. S1A, B**). Increased amounts of the mutated enzyme showed no improvement in tagmentation, verifying the important role of the DDE domain in Tn5 tagmentation. Tn5 is a member of the ribonuclease H-like (RNHL) superfamily, together with RNase H and MMLV reverse transcriptase (35, 39, 40); therefore, we hypothesized that Tn5 is capable of processing not only dsDNA but also RNA/DNA heteroduplex. Sequence alignment between these three proteins revealed a conserved domain (two Asps and one Glu) within their active sites, termed the RNase H-like domain (**Fig. 1A**). The two Asp residues (D97 and D188) in the Tn5 catalytic core were structurally similar to those of the other two enzymes (**Fig. 1B**). Moreover, divalent ions, which are important for stabilizing substrate and catalyzing reactions, also occupy similar positions in all three proteins (**Fig. 1B**) (39). We determined the nucleic acid substrate binding pocket of Tn5 according to charge distribution. We then docked double-stranded DNA and RNA/DNA heteroduplex into this predicted pocket and showed that the binding site had enough space for an RNA/DNA duplex (**Fig. S1C**). These structural similarities among Tn5, RNase H and MMLV reverse transcriptase and the docking results further supported the possibility that Tn5 can catalyze the strand transfer reaction on RNA/DNA heteroduplex (**Fig. 1C**).

**Fig. 1.**
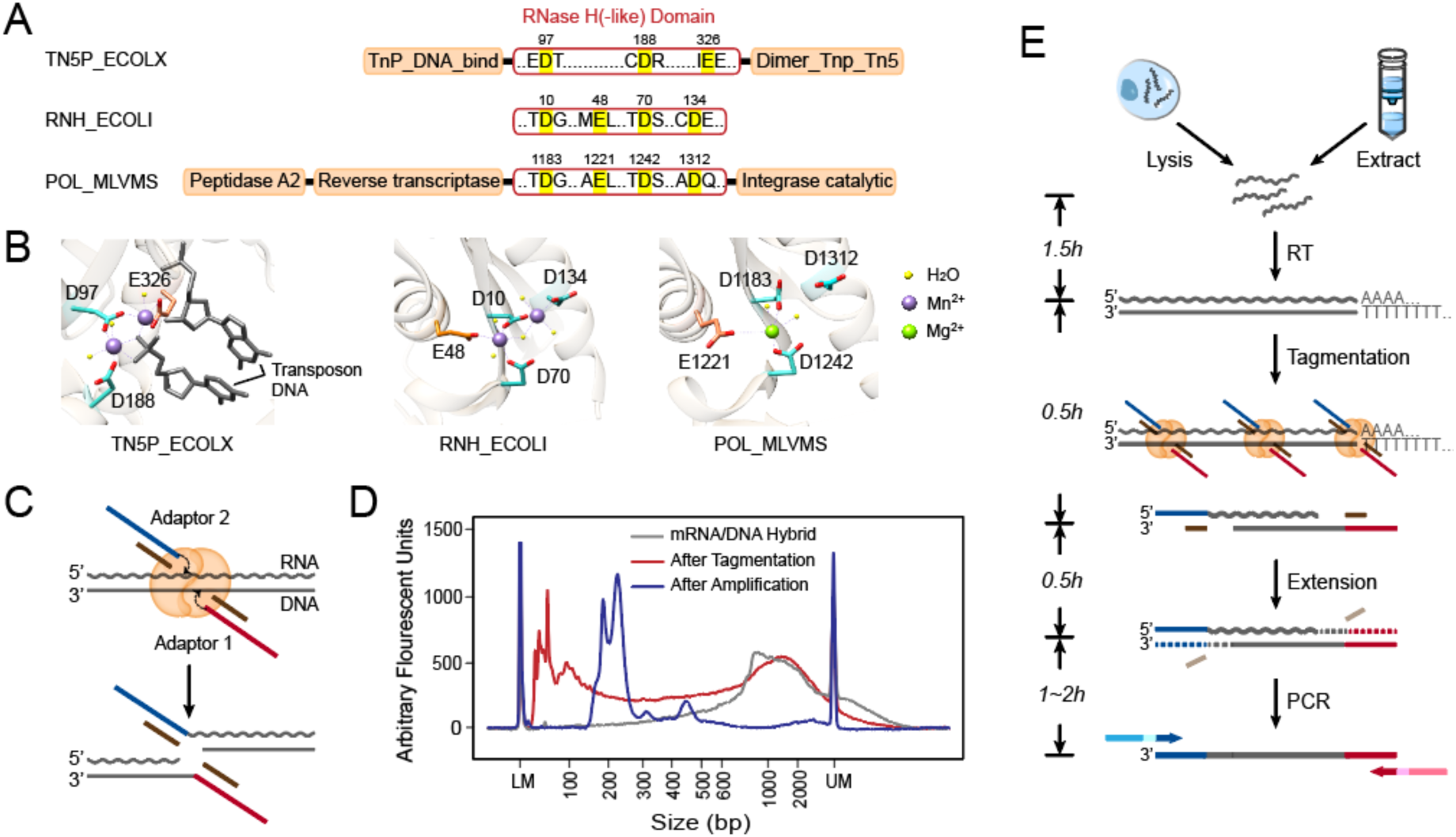
Tn5 tagmentation activity on double-stranded hybrids and the experimental process of SHERRY. (**A**) RNase H-like (RNHL) domain alignment of Tn5 (TN5P_ECOLX), RNase H (RNH_ECOLI) and M-MLV reverse transcriptase (POL_MLVMS). Active residues in the RNHL domains are labeled in bright yellow. Orange boxes represent other domains. (**B**) Superposition of the RNHL active sites in these three enzymes. PDB IDs are 1G15 (RNase H), 2HB5 (M-MLV) and 1MUS (Tn5). (**C**) Putative mechanism of Tn5 tagmentation of an RNA/DNA hybrid. Crooked arrows represent nucleophilic attacks. (**D**) Size distribution of mRNA/DNA hybrids with and without Tn5 tagmentation and after amplification with index primers. (**E**) Workflow of sequencing library preparation by SHERRY. The input can be a lysed single cell or extracted bulk RNA. After reverse transcription with oligo-dT primer, the hybrid is directly tagmented by Tn5, followed by gap-repair and enrichment PCR. Wavy and straight gray lines represent RNA and DNA, respectively. Dotted lines represent the track of extension step.

To validate our hypothesis, we purified Tn5 using the pTXB1 plasmid and corresponding protocol. We prepared RNA/DNA hybrids using mRNA extracted from HEK293T cells. Using a typical dsDNA tagmentation protocol, we treated 15 ng of RNA/DNA hybrids with 0.6 µl pTXB1 Tn5 transposome. Fragment analysis of the tagmented RNA/DNA hybrids showed an obvious decrease (∼1000bp) in fragment size compared with that of untreated control, validating the capability of Tn5 to fragment the hybrid (**Fig. 1D**).

Based on the ability of the Tn5 transposome to fragment RNA/DNA heteroduplexes, we propose SHERRY (Sequencing HEteRo RNA-DNA-hYbrid), a rapid RNA-seq library construction method (**Fig. 1E**). SHERRY consists of three components: RNA reverse transcription, RNA/cDNA hybrid tagmentation, and PCR amplification. The resulting product is an indexed library that is ready for sequencing. Specifically, mRNA is reverse transcribed into RNA/cDNA hybrids using d(T)_30_VN primer. The hybrid is then tagmented by the pTXB1 Tn5 transposome, the adding of partial sequencing adaptors to fragment ends. DNA polymerase then amplifies the cDNA into a sequencing library after initial end extension. The whole workflow only takes approximately four hours with hand-on time less than 30 minutes.

To test SHERRY feasibility, we gap-repaired the RNA/DNA tagmentation products illustrated in **Fig. 1D** (red line) and then amplified these fragments with library construction primers. Amplified molecules (**Fig. 1D**, blue line) were approximately 100∼150 bp longer than the tagmentation products, which matched the extra length of adaptors added by gap-repair and index primer amplification. (**Fig. S2**). Thus, direct Tn5 tagmentation of RNA/DNA hybrids offers a new strategy for RNA-seq library preparation.

### Tn5 has ligation activity on tagmented RNA/DNA hybrids

To further investigate the detailed molecular events of RNA/DNA hybrid tagmentation, we designed a series of verification experiments. First, we wanted to verify that the transposon adaptor can be ligated to the end of fragmented RNA (**Fig. 2A**). In brief, we prepared RNA/DNA hybrids from HEK293T RNA by reverse transcription. After tagmentation with the Tn5 transposome, we purified the products to remove Tn5 proteins and free adaptors. We assumed that Tn5 ligated the adaptor to the fragmented DNA. At the same time, if Tn5 ligated the adaptor (**Fig. 2A**, dark blue) to the RNA strand, the adaptor could serve as a template in the subsequent extension step. After extension the DNA strand should have a primer binding site on both 5′ and 3′ ends for PCR amplification. RNase H treatment should not affect production of the sequencing library. If Tn5 failed to ligate the adaptor to the RNA strand, neither strand of the heteroduplex would be converted into a sequencing library.

**Fig. 2.**
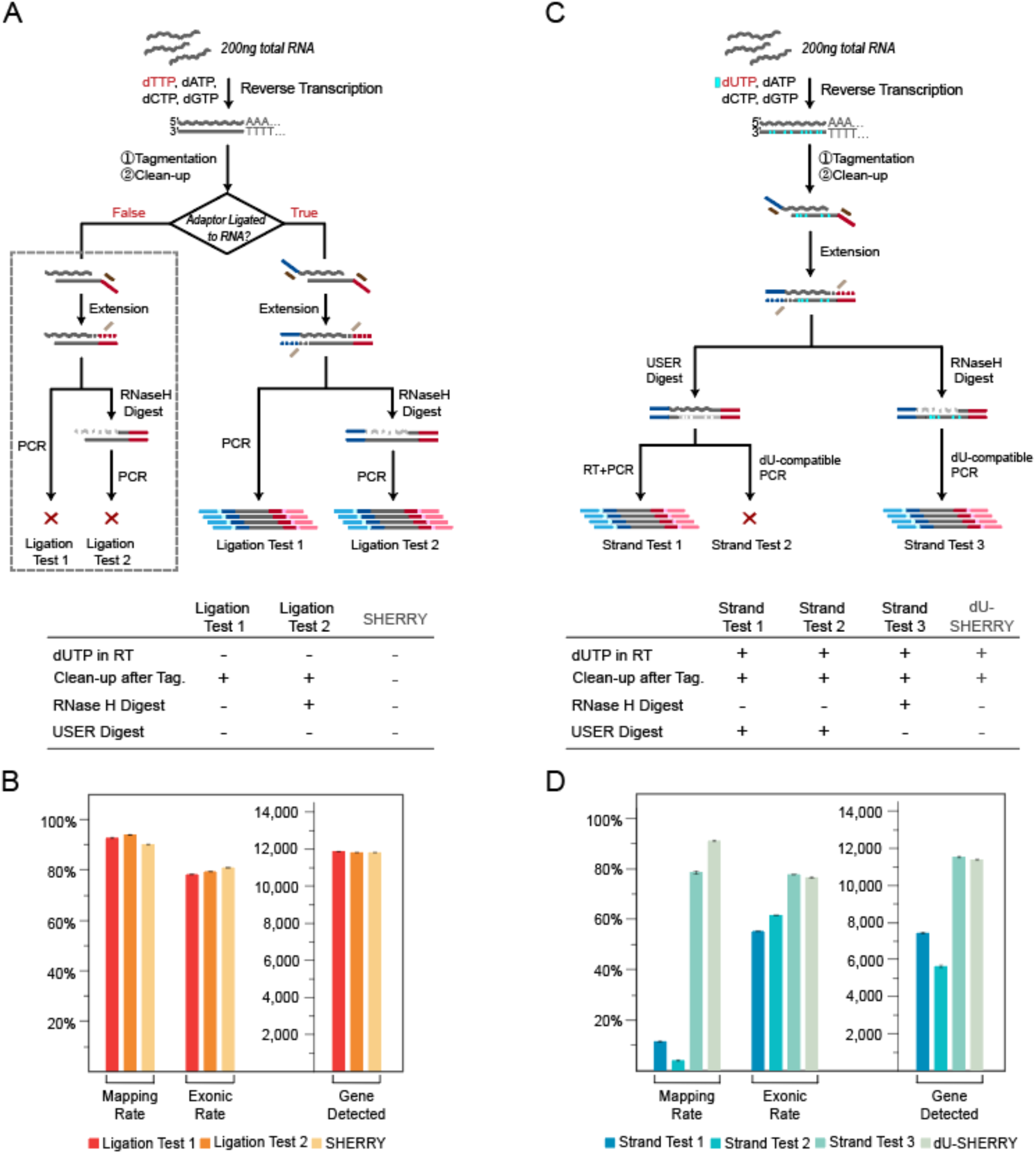
Verification of Tn5 tagmentation of RNA/DNA heteroduplexes. (**A**) Procedures of two Ligation Tests. Gray dotted box indicates negative results. The table below lists key experimental parameters that are different from standard SHERRY. (**B**) Comparison of two Ligation Tests and standard SHERRY with respect to mapping rate, exon rate and number of genes detected. Each test consisted of two replicates of 200 ng HEK293T total RNA. (**C**) Strand Test procedures. (**D**) Comparison among three Strand Tests and dU-SHERRY with respect to mapping rate, exon rate and number of genes detected. Each test consisted of two replicates of 200 ng HEK293T total RNA.

After PCR amplification, we obtained a high quantity product regardless of RNase H digestion, indicating successful ligation of the adaptor to the fragmented DNA. Sequencing results from both reaction test conditions as well as from SHERRY showed >90% mapping rate to the human genome with ∼80% exon rate and nearly 12,000 genes detected, validating the transcriptome origin of the library (**Fig. 2B, Fig. S3A**). The additional purification step after reverse transcription and/or RNase H digestion before PCR amplification did not affect the results, probably because of the large amount of starting RNA. We examined the sequencing reads with an insert size shorter than 100 bp (shorter than the sequence read length), and 99.7% of them contained adaptor sequence. Such read-through reads directly proved ligation of the adaptor to the fragmented RNA (**Fig. S3B and C**). In summary, we confirmed that Tn5 transposome can tagment both DNA and RNA strands of RNA/DNA heteroduplexes.

### Tagmented cDNA is the preferred amplification template

Next, we investigated whether RNA and DNA strands could be amplified to form the sequencing library (**Fig. 2C**). We replaced dTTP with dUTP during the reverse transcription and then purified the tagmented products to remove free dUTP and Tn5 proteins. Bst 2.0 Warmstart DNA polymerase was used for extension because it is able to use RNA as a primer and to process the dU bases in the template. The product fragments were then treated with either USER enzyme or RNase H to digest cDNA and RNA, respectively. We performed RT-PCR with the USER-digested product, to test the efficiency of converting tagmented RNA for library construction (Strand Test 1). To exclude interference from undigested DNA, we performed PCR amplification with the USER-digested fragments using dU-compatible polymerase (Strand Test 2). We also used dU-compatible PCR to test the efficiency of converting tagmented cDNA for library construction (Strand Test 3). For comparison, we included a control experiment with the same workflow as Strand Test 3 except that the RNase H digestion step was omitted (dU-SHERRY) to ensure that Tn5 can recognize substrates with dUTP.

Sequencing results of Strand Test 1 showed a low mapping rate and gene detection count that were only slightly higher than those of Strand Test 2. In contrast, Strand Test 3 demonstrated a similar exon rate and gene count to dU-SHERRY and SHERRY (**Fig. 2D, Fig. S3A**). Based on these results, we conclude that the tagmented cDNA contributes to the majority of the final sequencing library, likely because of inevitable RNA degradation during the series of reactions.

### SHERRY for rapid one-step RNA-seq library preparation

We tested different reaction conditions to optimize SHERRY with 10 ng total RNA as input (**Fig. S4A**). We evaluated the impact of different crowding agents, different ribonucleotide modifications on transposon adaptors, and different enzymes for gap filling. We also included purification after certain steps to remove primer dimers and carry-over contaminations. Sequencing results showed little change in performance from most of these modifications, indicating that SHERRY is robust under various conditions.

We then compared the optimized SHERRY with NEBNext® Ultra™ II, a commercially available kit, for bulk RNA library preparation. This NEBNext kit is one of the most commonly used kits used for RNA-seq experiments, with 10 ng total RNA being its minimum input limit. We therefore tested the RNA-seq performance with 10 ng and 200 ng total RNA inputs, each condition having three replicates. SHERRY demonstrated comparable performances with NEBNext for both input levels (**Fig. S4B**). For the 10 ng input tests, SHERRY produced more precise gene expression measurements across replicates (**Fig. 3A**), probably because of the simpler SHERRY workflow.

**Fig. 3.**
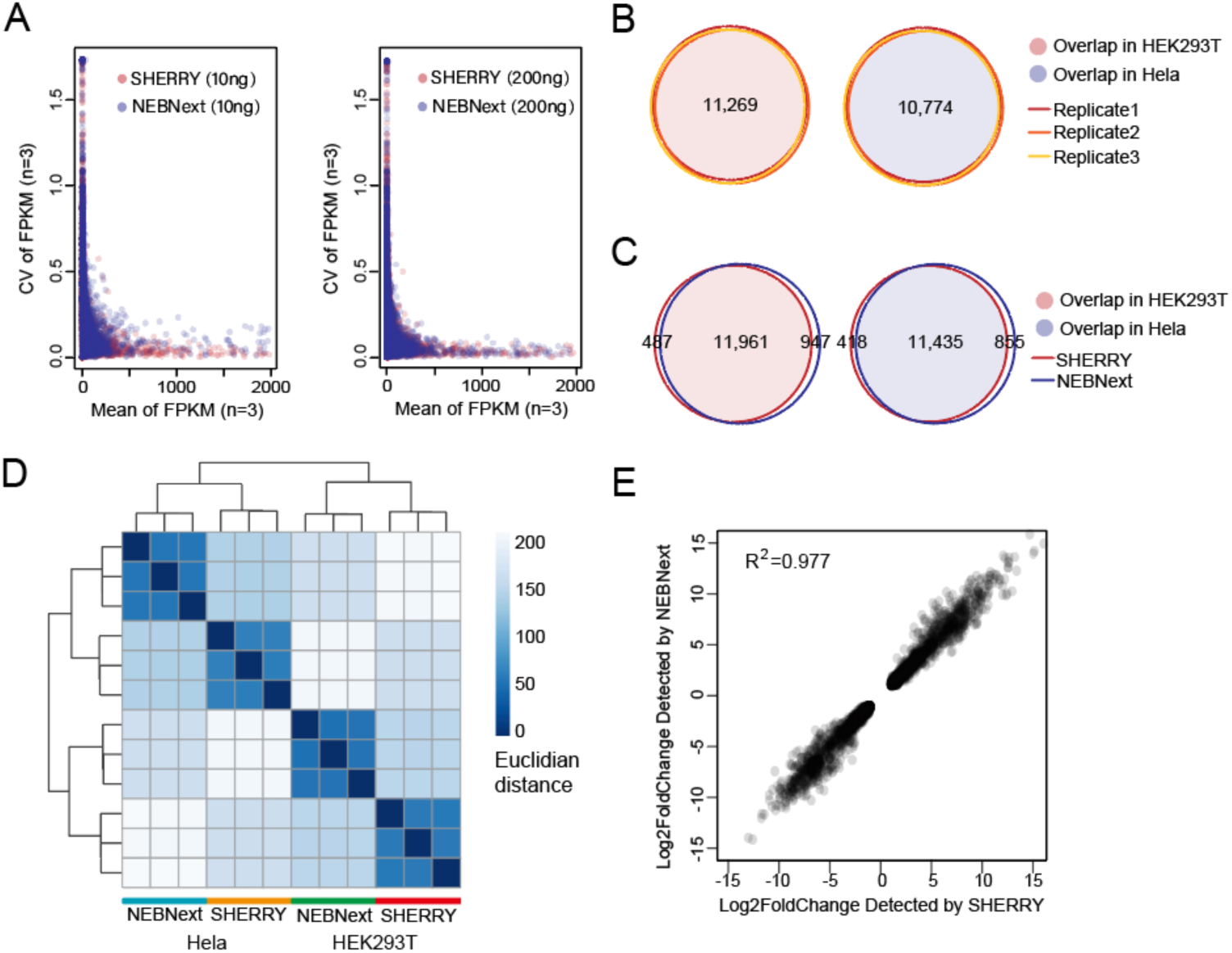
Performance of SHERRY with large RNA input. (**A**) Coefficient of variation (CV) across three replicates was plotted against the mean value of each gene’s FPKM (Fragments Per Kilobase of transcript per Million mapped reads). All experiments used HEK293T total RNA as input. (**B**) Genes detected by SHERRY in three replicates of 200 ng HEK293T or HeLa total RNA are plotted in Venn Diagrams. Numbers of common genes are indicated. (**C**) Common genes detected by SHERRY and NEBNext in the three replicates of 200 ng HEK293T or HeLa total RNA. (**D**) Distance heatmap of samples prepared by SHERRY or NEBNext for three replicates using 200 ng HEK293T or HeLa total RNA. The color bar indicates the Euclidian distance. (**E**) Correlation of gene expression fold-change identified by SHERRY and NEBNext. Involved genes are differentially expressed genes between HEK293T and HeLa detected by both methods.

Next, we compared the ability to detect differentially expressed genes between HEK293T and HeLa cells using SHERRY and NEBNext. In all three replicates, SHERRY detected 11,269 genes in HEK293T cells and 10,774 genes in HeLa cells, with high precision (correlation coefficient R^2^=0.999) (**Fig. 3B, Fig. S5A**). The numbers of detected genes and their read counts identified by SHERRY and NEBNext were highly concordant (**Fig. 3C, Fig.S5B**). This excellent reproducibility of SHERRY ensured the reliability of subsequent analyses. Then we plotted a heatmap of the distance matrix (**Fig. 3D**) between different cell types and library preparation methods. Libraries from the same cell type were clustered together as expected. Libraries from the same method also tended to cluster together, indicating internal bias in both methods.

We then used DESeq2 to detect differentially expressed genes (P-value <5×10^−6^, |log_2_Fold change| >1). In general, the thousands of differentially expressed genes detected by both methods were highly similar (**Fig. S5C**) and their expression fold-change was highly correlated (correlation coefficient R^2^=0.977) between SHERRY and NEBNext (**Fig. 3E**). Examination of the genes that showed differential expression in only one method, revealed the same trend of expression change in the data from the other method (**Fig. S5D and E**). We conclude that SHERRY provides equally reliable differential gene expression information as NEBNext, but with a much faster and less labor-intensive process, specifically saving around two hours hand-on time (**Fig. S7A**).

### SHERRY using trace amounts of RNA or single cells

We next investigated whether SHERRY could construct RNA-seq libraries from single cells. First, we reduced the input to 100 pg total RNA, which is equivalent to RNA from about 10 cells. SHERRY results were high quality, with high mapping and exon rates and nearly 9,000 genes detected (**Fig. S6A**). 72% of these genes were detected in all three replicates, demonstrating good reproducibility (**Fig. 4A**). The expression of these genes showed excellent precision with R^2^ ranging from 0.958 to 0.970. (**Fig. 4B**).

**Fig. 4.**
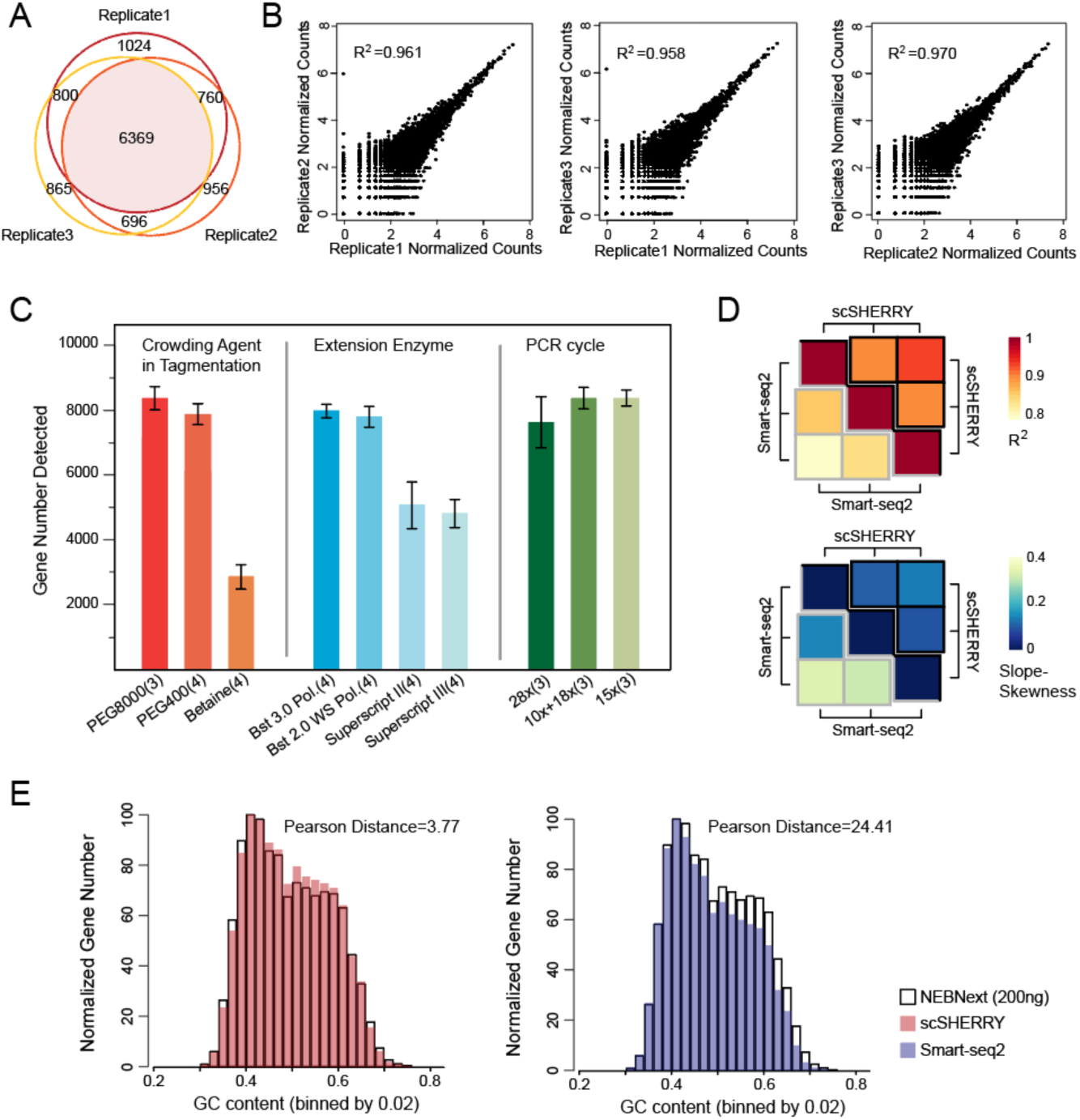
Performance of SHERRY with micro-input samples. (**A**) Genes detected by SHERRY in three replicates with 100 pg HEK293T total RNA. (**B**) Correlations of normalized gene read counts between replicates with 100 pg HEK293T total RNA. (**C**) Gene number detected by scSHERRY under various experimental conditions in single HEK293T cells. Each condition involved 3–4 replicates. (**D**) The heatmap of R^2^ calculated from scSHERRY and Smart-seq2 replicates, and slope deviation in a linear fitting equation for the two methods. (**E**) Normalized gene numbers with different GC content.

To further push the detection limit, we carried out single-cell SHERRY experiments (scSHERRY) using HEK293T cell line. In contrast to the experiments with purified RNA, scSHERRY required several optimizations to the standard protocol (**Fig. 4C, Fig. S6B**). Although we found no positive effect by replacing betaine in the standard protocol with other crowding agents during optimization, we found that addition of a crowding reagent with a higher molecular weight improved the library quality from single cells. Therefore, we used PEG8000 for the following scSHERRY experiments. For the extension step, the use of Bst 3.0 or Bst 2.0 WarmStart DNA polymerases detected more genes than the use of Superscript II or Superscript III reverse transcriptases. This is probably because of the stronger processivity and strand displacement activity of Bst polymerases, and better compatibility with higher reaction temperatures to open the secondary structure of RNA templates. We also tried to optimize the PCR strategy because extensive amplification can lead to strong bias. Compared to the continuous 28-cycle PCR, the incorporation of a purification step after 10 cycles, or simply reducing the total cycle number to 15 increased the mapping rate and the number of genes detected. Therefore, we performed 15-cycle PCR without extra purification to better accommodate high-throughput experiments.

The optimized scSHERRY was capable of detecting 8,338 genes with a 50.17% mapping rate (**Fig. S6D**), and the gene read counts correlated well (correlation coefficient R^2^=0.600) with Smart-seq2, the most prevalent protocol in the single-cell RNA amplification field. Besides, scSHERRY showed better reproducibility than Smart-seq2 (**Fig.4D**). Compared with Smart-seq2, the gene number and coverage uniformity of the scSHERRY-generated library was slightly inferior (**Fig. S6D, E**) because Smart-seq2 enriches for full-length transcripts via a preamplification step. However, this enrichment step of Smart-seq2 also introduced bias (**Fig. 4E**). We used 200 ng of HEK293T total RNA to construct a sequencing library using the NEBNext kit, expecting to capture as many genes as possible. Besides, the NEBNext protocol used short RNA fragments for reverse transcription and fewer cycles of PCR amplification, which should introduce less GC bias than the other protocols. We then compared the GC distribution of genes detected by scSHERRY or Smart-seq2 with NEBNext results. scSHERRY, which is free from second-strand synthesis and preamplification, produced a distribution similar to the standard. However, the library from Smart-seq2-amplified scRNA showed clear enrichment for genes with lower GC content. Genes with high GC content were less likely to be captured by Smart-seq2, which may cause biased quantitation results. Overall, compared with Smart-seq2, scSHERRY produced libraries of comparable quality and lower GC-bias. Moreover, the scSHERRY workflow spares pre-amplification and QC steps before tagmentation, saving around four hours (**Fig. S7A**), and the one tube strategy is promising for high throughput application.

## Discussion

We found that the Tn5 transposome has the capability to directly fragment and tag RNA/DNA heteroduplexes and, therefore, we have developed a quick RNA amplification and library preparation method called SHERRY. The input for SHERRY could be RNA from single cell lysate or total RNA extracted from a large number of cells. Comparison of SHERRY with the commonly used Smart-seq2 protocol for single-cell input or the NEBNext kit for bulk total RNA input, showed comparable performance for input amount spanning more than five orders of magnitude. Furthermore, the whole SHERRY workflow from RNA to sequencing library consists of only five steps in one tube and takes about four hours, with hands-on time of less than 30 min. Smart-seq2, requires twice this amount of time and an additional library preparation step is necessary. The ten-step NEBNext protocol is much more labor-intensive and time-consuming (**Fig. S7A**). Moreover, the SHERRY reagent cost is five-fold less compared with that of the other two methods (**Fig. S7B**). For single cells, the lower mapping rate of SHERRY compared to Smart-seq2 could increase sequencing cost. However, both methods reached a plateau of saturation curve with 2 million total reads (**Fig. S8**), which costs less than $5. Therefore, SHERRY has strong competitive advantages over conventional RNA library preparation methods and scRNA amplification methods.

In our previous experiments, we assembled a Tn5 transposome using home-purified pTXB1 Tn5 and synthesized sequencer-adapted oligos. To generalize the SHERRY method, and to confirm Tn5 tagmentation of RNA/DNA heteroduplexes, we tested two commercially available Tn5 transposomes, Amplicon Tagment Mix (abbreviated as ATM) from the Nextera XT kit (Illumina, USA) and TTE Mix V50 (abbreviated as V50) from the TruPrep kit (Vazyme, China). We normalized the different Tn5 transposome sources according to the enzyme processing 5 ng of genomic DNA to the same size under the same reaction conditions (**Fig. S9A**). The tagmentation activity of our in-house pTXB1 Tn5 was 10-fold higher than V50 and 500-fold higher than ATM when using transposome volume as the metric. The same units of enzyme were then used to process RNA/DNA heteroduplexes prepared from 5 ng mRNA to confirm similar performance on such hybrids (**Fig. S9B**). The RNA-seq libraries from all three enzymes showed consistent results, demonstrating the robustness of SHERRY (**Fig. S9C**).

During DNA and RNA/DNA heteroduplex tagmentation, the Tn5 transposome reacted with these two substrates in different patterns. We tagmented 5 ng DNA or mRNA/DNA hybrids with 0.02 µl, 0.05 µl or 0.2 µl pTXB1 Tn5 transposome (**Fig. S10**). As the amount of Tn5 increased, dsDNA was cut into overall shorter fragments. While for the hybrid, Tn5 cut the template ‘one by one’, because only hybrids above a certain size became shorter and most were too short to be cut. We supposed that such phenomenon might attribute to the different conformation of dsDNA and RNA/DNA hybrid, since diameter of the latter is larger. Thus, the binding pocket or catalytic site of Tn5 would be tuned to accommodate the hybrid strands and cause different tagmentation pattern.

Despite its ease-of-use and commercial promise, the library quality produced by SHERRY may be limited by unevenness of transcript coverage. Unlike the NEBNext kit, which fragments RNA before reverse transcription, or Smart-seq2, which performs pre-amplification to enrich full-length cDNAs, SHERRY simply reverse transcribes full-length RNA. Reverse transcriptase is well known for its low efficiency and when using polyT as the primer for extension, it is difficult for the transcriptase to reach the 5′ end of the RNA template. This can cause coverage imbalance across transcripts, making the RNA-seq signal biased toward the 3′ end of genes (**Fig. S11A**). In an attempt to solve this problem, we added TSO primer, the sequence and concentration of which was the same as Smart-seq2 protocol (17), to the reverse transcription buffer to mimic the Smart-seq2 reverse transcription conditions. The resulting hybrid was then tagmented and amplified following standard SHERRY workflow. This produced much improved evenness across transcripts (**Fig. S11A and B**), although some of the sequencing parameters dropped accordingly (**Fig. S11C**). We believe that with continued optimization SHERRY will improve RNA-seq performance.

## Author contributions

Y.F., K.L., X.S.X., Y.H. and J.W. conceived the project; L.D., J.X., J.O. and D.W. performed structural analysis; Y.S., J.L. and G.W. performed Tn5 purification and characterization; L.D., Y.F. and L.L. conducted research; L.D., Y.F., L.L., J.Y., Y.W., R.L., G.Z., Y.H. and J.W. analyzed the data; L.D., X.S.X., Y.H. and J.W. wrote the manuscript with the help from all other authors.

## Conflict of interest statement

XGen US Co has applied for a patent related to this work. X. Sunney Xie, Kaiqin. Lao and Yalei Wu are shareholders of XGen US Co. Kai Q. Lao, Yalei Wu, Raymond W. Lee and Genhua Zheng are employees of XGen US Co.

## Acknowledgement

We thank Dr. Yun Zhang and BIOPIC sequencing platform at Peking University for the assistance of high-throughput sequencing experiments. This work was supported by National Natural Science Foundation of China (21675098, 21525521), Ministry of Science and Technology of China (2018YFA0800200, 2018YFA0108100, 2018YFC1002300), 2018 Beijing Brain Initiation (Z181100001518004), Beijing Advanced Innovation Center for Structural Biology, and Beijing Advanced Innovation Center for Genomics.

## Supplementary Figures

**Fig. S1.**
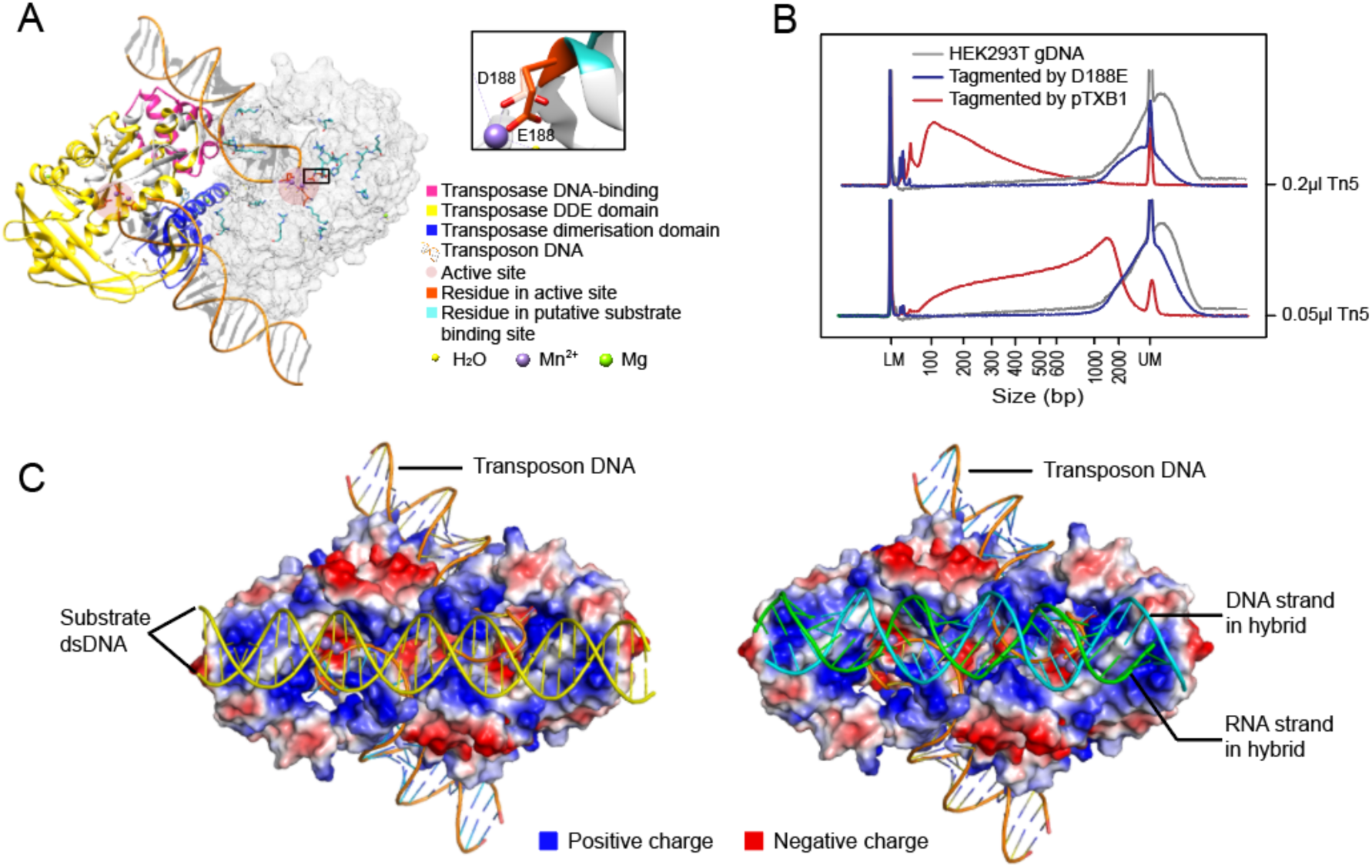
Structure of Tn5 and D188E mutation in Tn5. (**A**) Structure of pTXB1 Tn5 (PDB ID: 1MUS). Left monomer marked domains in different colors. Right monomer marked residues of catalytical core and putative substrate binding site in atom form. [Referred to D. R. Davies, I. Y. Goryshin, W. S. Reznikoff, I. Rayment, Three-dimensional structure of the Tn5 synaptic complex transposition intermediate. Science 289, 77-85 (2000).] Black box showed D188E mutation. (**B**) Size distribution of genomic DNA with no treatment or tagmented by D188E mutant Tn5 or tagmented by pTXB1 Tn5. (**C**) Model of docking double-stranded DNA (Left) or RNA/DNA heteroduplex (Right) in predicted substrate binding sites of Tn5.

**Fig. S2.**
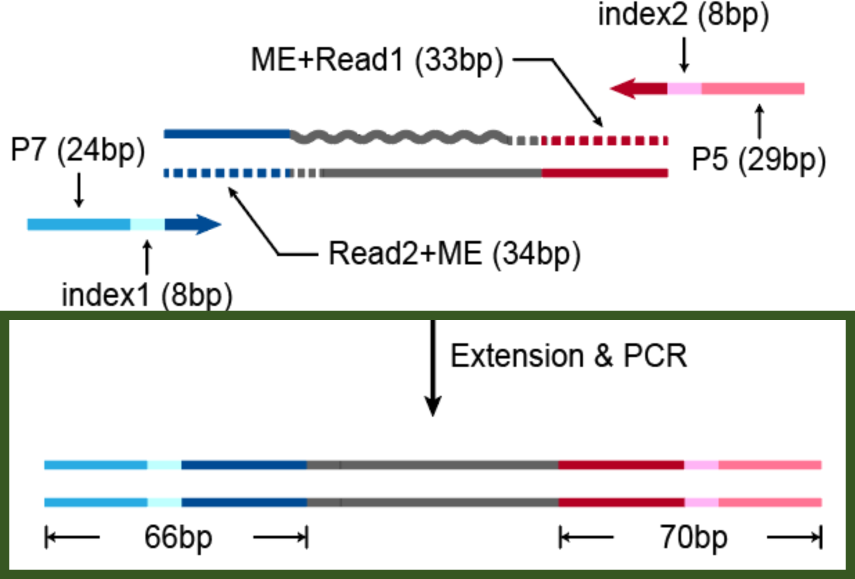
Composition of products amplified from tagmented RNA/DNA hybrid. Gray wavy line and straight line represent RNA and DNA separately. Dotted lines represent the track of extension step.

**Fig. S3.**
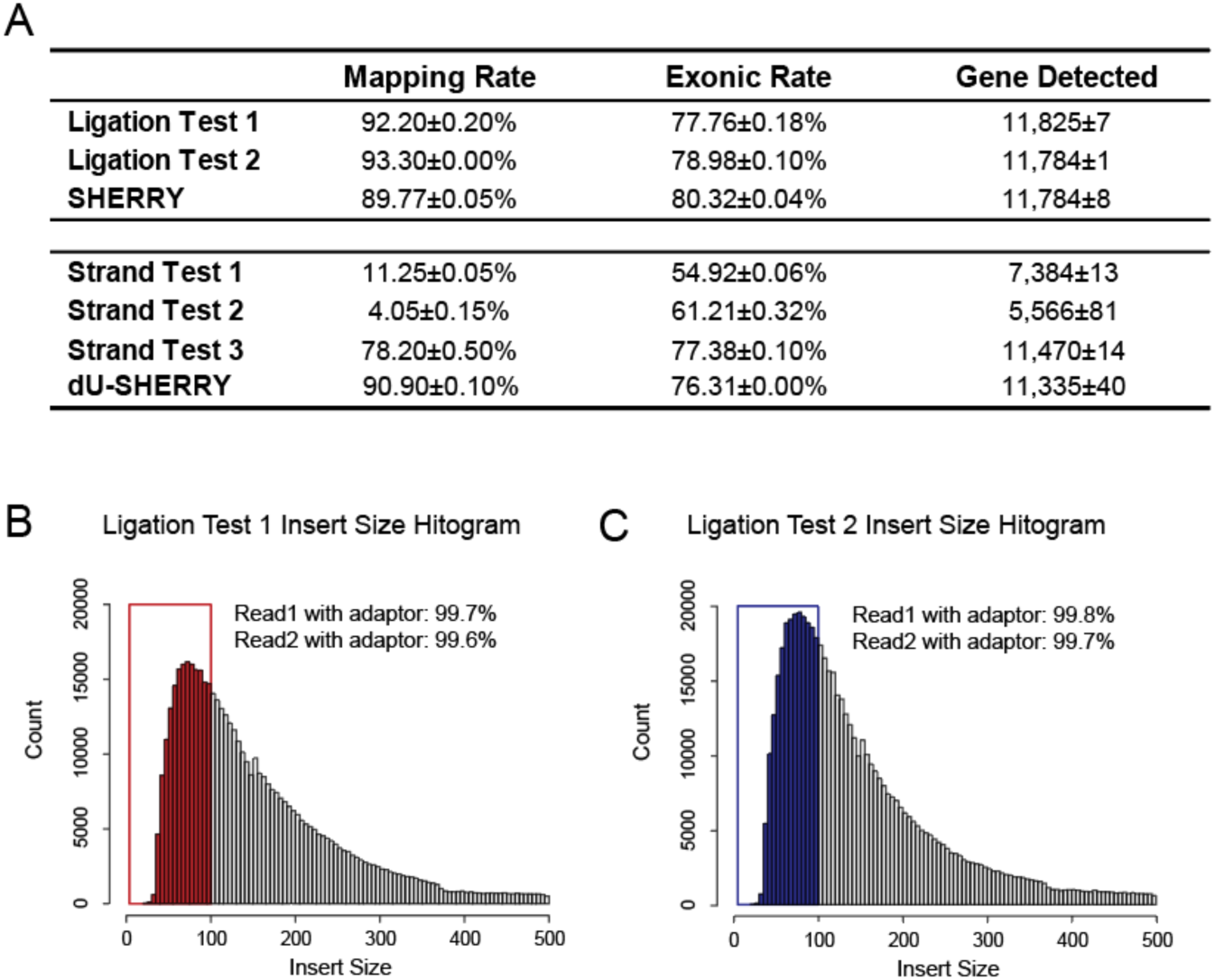
Comparison within Ligation Tests and Strand Tests. (**A**) Sequencing indicators of Ligation Tests and Strand Tests. Each test consisted of two replicates of 200 ng HEK293T total RNA. (**B-C**) Insert size distribution of Ligation Tests. The colored bars marked reads which insert size is shorter than 100bp. Adaptors detected in these reads are counted.

**Fig. S4.**
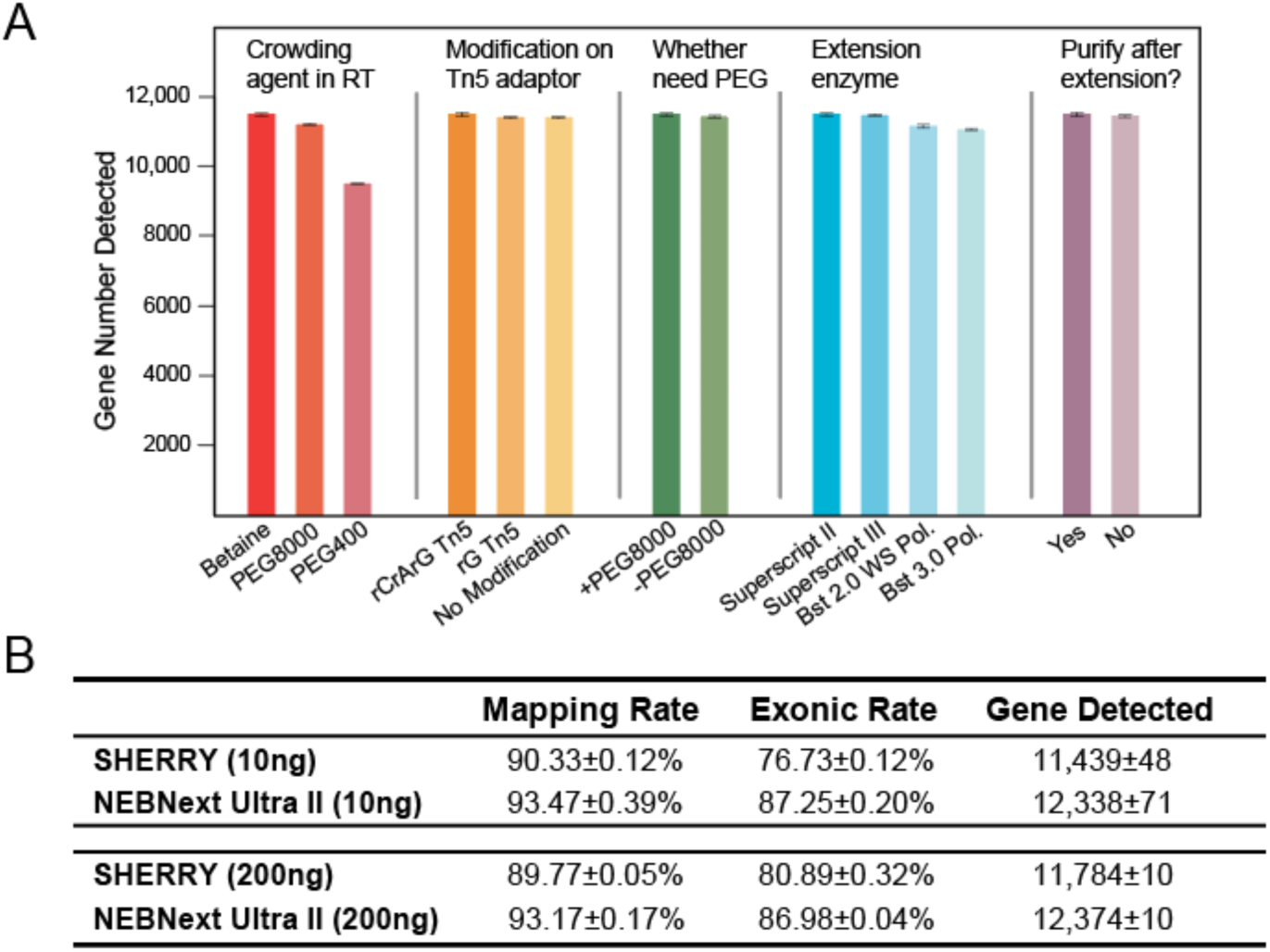
Optimization of SHERRY and comparison with NEBNext. (**A**) Gene number detected by SHERRY under various experimental conditions. Each condition consisted of three replicates of 10 ng HEK293T total RNA. (**B**) Comparison of sequencing indicators between SHERRY and NEBNext with 10 ng and 200 ng HEK293T total RNA input. Each condition consisted of three replicates and down-sampled to 2 million total reads.

**Fig. S5.**
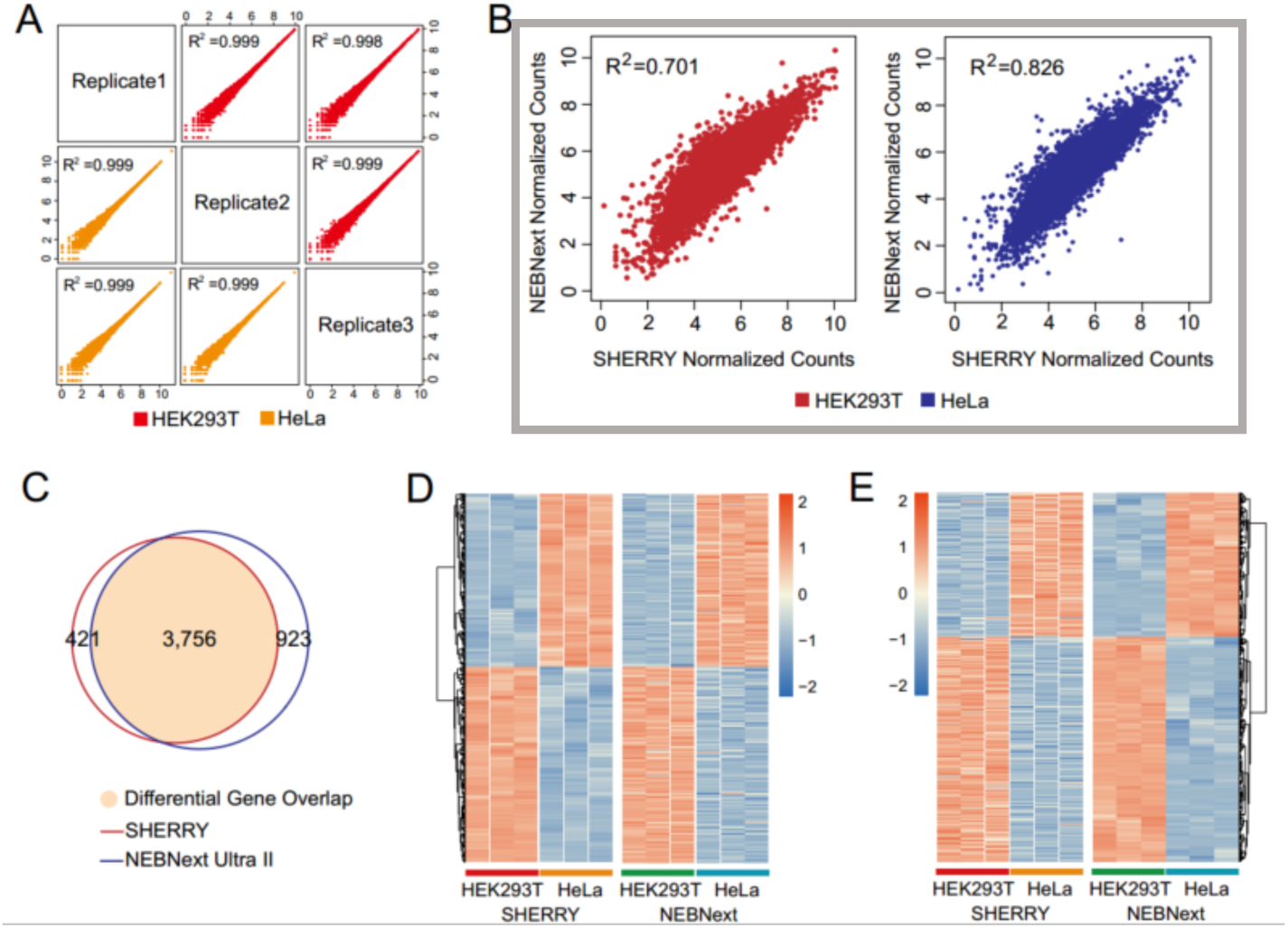
Functional comparison between SHERRY and NEBNext. (**A**) Correlation of normalized gene counts among duplicates of SHERRY, which start from 200 ng HEK293T total RNA input. (**B**) Correlation of normalized genes counts (average of three replicates) between SHERRY and NEBNext within the two cell types. The input was 200 ng total RNA. (**C**) Differentially expressed genes of HeLa and HEK293T detected by SHERRY and NEBNext kit (200 ng input) are plotted into Venn Diagram. Colored area represents genes identified by both methods. Gene numbers are listed on corresponding part. (**D**) Heatmap of differentially expressed genes detected by SHERRY while missed by NEBNext kit. The Color bar indicates Z-score. (**E**) Heatmap of differentially expressed genes detected by NEBNext kit while missed by SHERRY. The Color bar indicates Z-score.

**Fig. S6.**
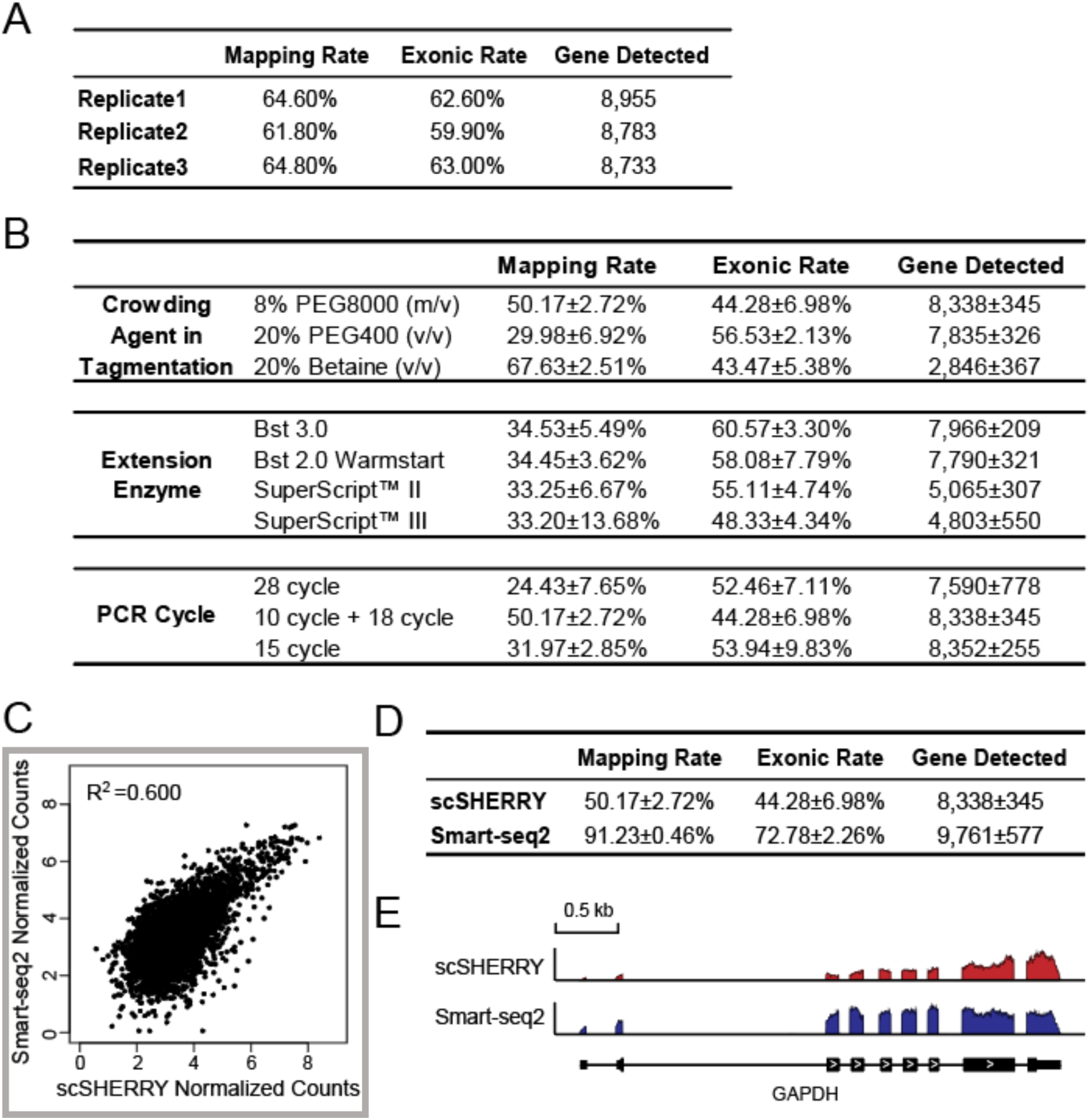
Optimization of micro-input SHERRY and comparison with Smart-seq2. (**A**) Sequencing indicators of SHERRY (n=3) starting from 100 pg HEK293T total RNA input. (**B**) Comparison of scSHERRY library quality under various experiment conditions. Each condition used single HEK293T cell as input and consisted of 3-4 replicates. (**C**) Correlation of normalized gene counts (average of three replicates) between scSHERRY and Smart-seq2. (**D**) Comparison of sequencing indicators between scSHERRY (n=3) and Smart-seq2 (n=4). Both of them used single HEK293T cells as input. (**E**) The coverage of GAPDH transcript calculated from scSHERRY and Smart-seq2.

**Fig. S7.**
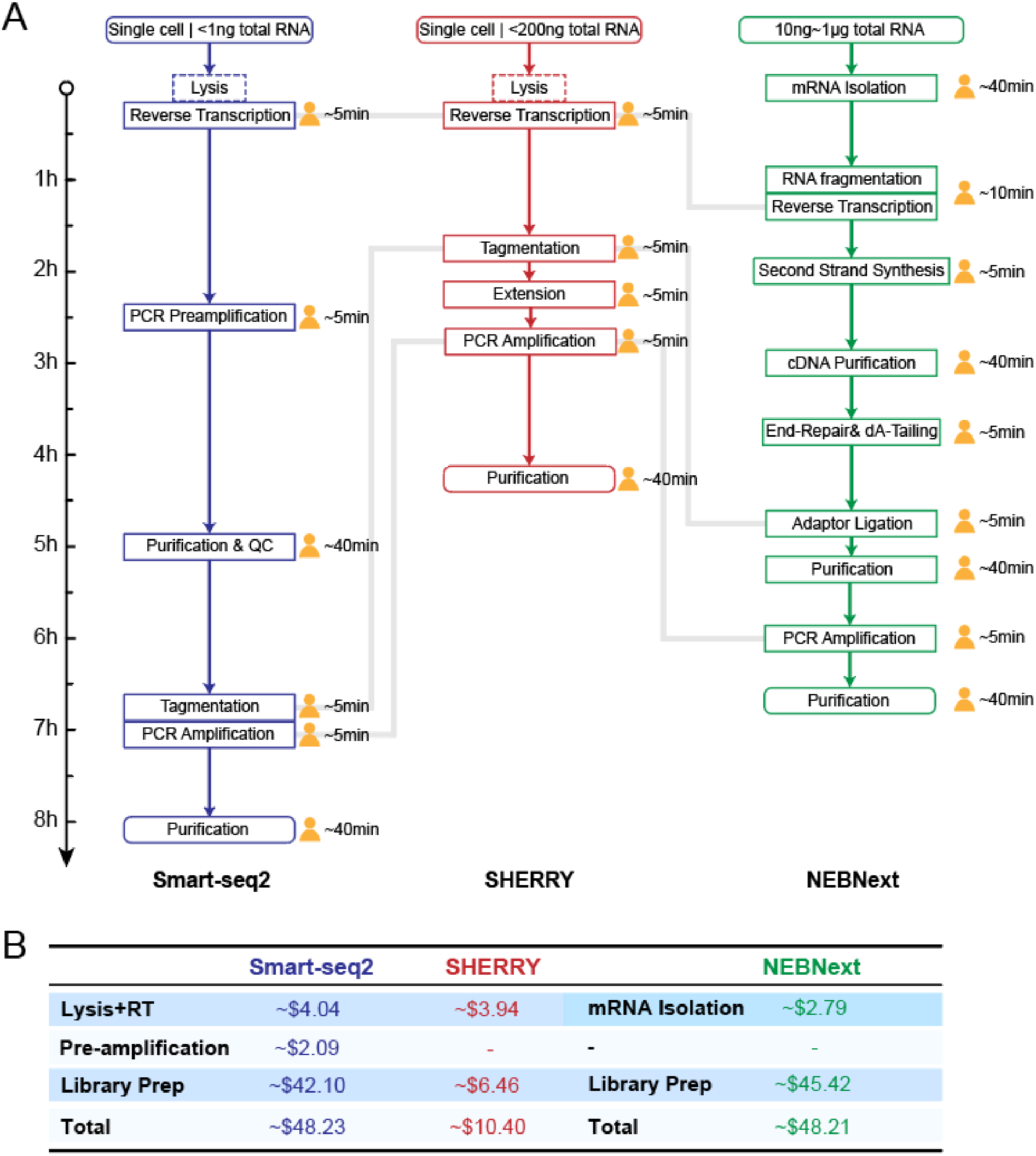
Comparison of workflow and cost among Smart-seq2, SHERRY and NEBNext kit. **(A)** Workflow of Smart-seq2, SHERRY and NEBNext kit. Length of arrow indicates time consumed for each step. The human-shaped icon indicated hands-on time. Dotted box means this step is alternative. Gray line connects corresponding key step in each method. (**B**) Cost list of Smart-seq2, SHERRY and NEBNext kit.

**Fig. S8.**
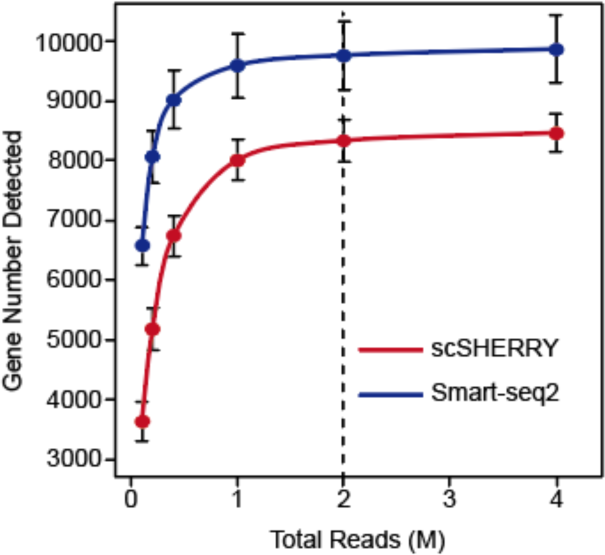
Saturation curve of scSHERRY (n=3) and Smart-seq2 (n=4). Both methods used single HEK293T cell as input. Dotted line indicated 2 million total reads.

**Fig. S9.**
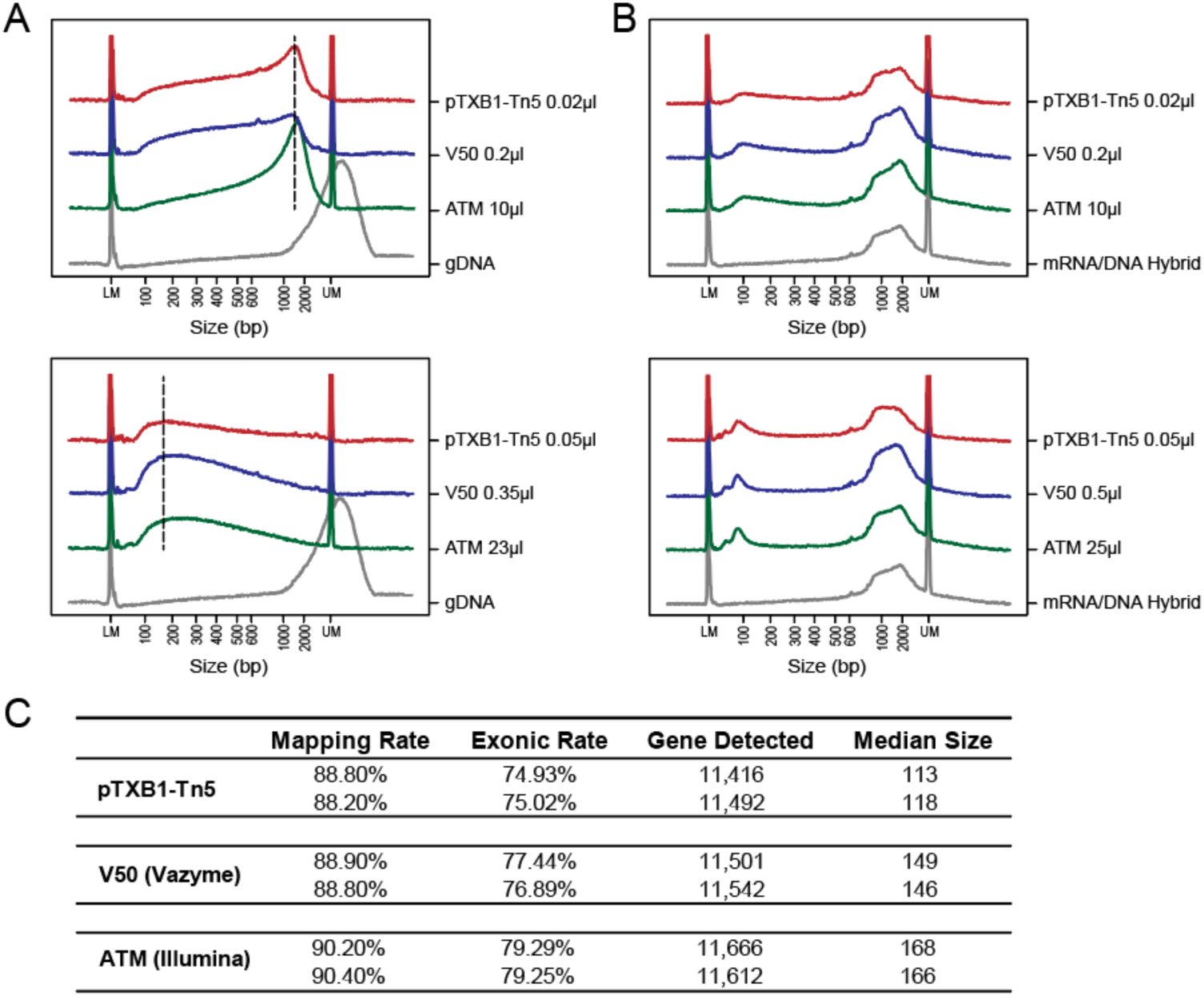
Hybrid tagmentation activity of commercial Tn5. (**A**) Size distribution of genomic DNA with no treatment or tagmented by different volumes of pTXB1 Tn5, V50 or ATM. The dotted black line indicates peak of fragment size. (**B**) Size distribution of mRNA/DNA hybrid with no treatment or tagmented by different volumes of pTXB1 Tn5, V50 or ATM. (**C**) Sequencing indicators of SHERRY library constructed by three Tn5.

**Fig. S10.**
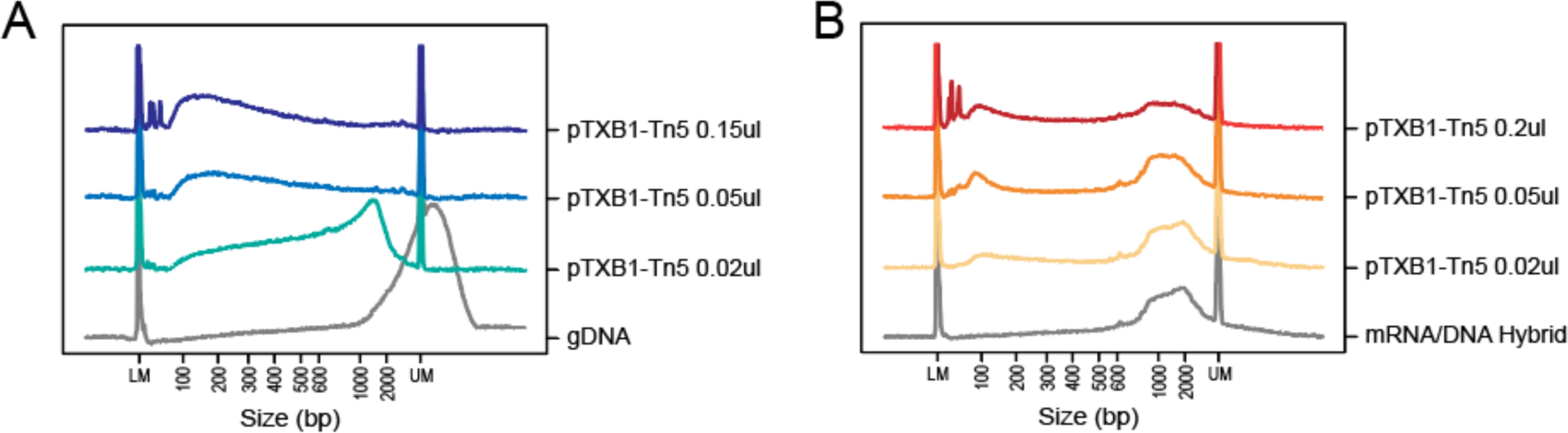
Titration of Tn5 transposome for tagmentation. (**A-B**) Size distribution of 5ng genomic DNA or mRNA/DNA hybrid with no treatment or tagmented by different gradients of pTXB1 Tn5.

**Fig. S11.**
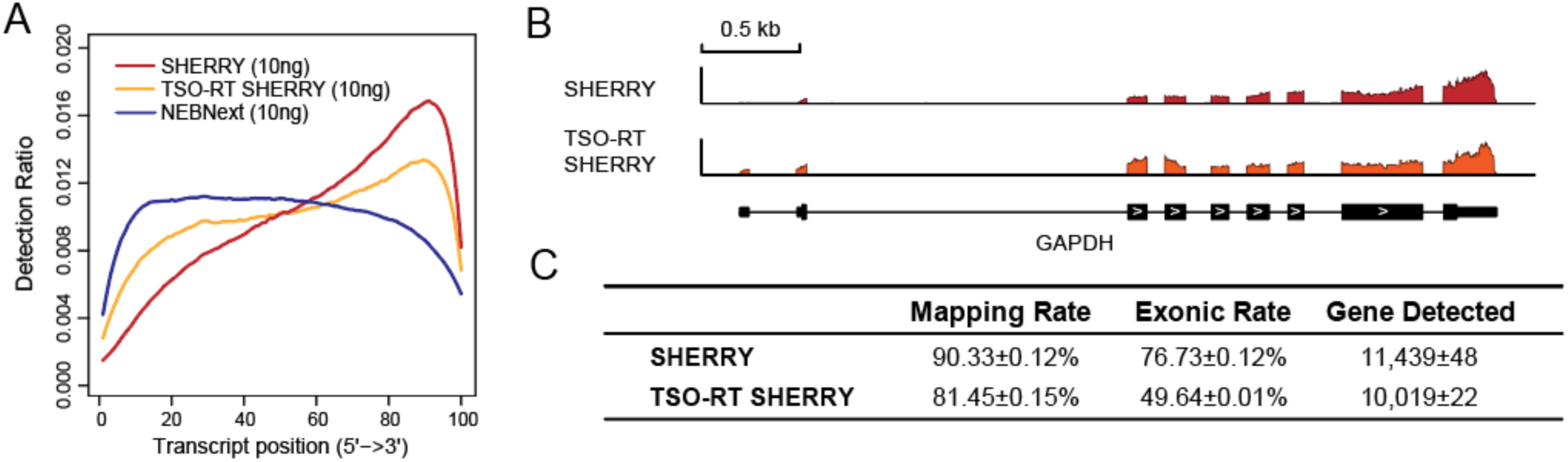
Coverage evenness optimization of SHERRY (10 ng HEK293T total RNA input). (**A**) Normalized transcript coverage of standard SHERRY, SHERRY using TSO-RT method and NEBNext kit. (**B**) The coverage of GAPDH transcript calculated from SHERRY and TSO-RT SHERRY. (**C**) Comparison of sequencing indicators between SHERRY (n=3) and TSO-RT SHERRY (n=2).

## Materials and Methods

### Purification of pTXB1 Tn5 and D188E mutation

The pTXB1 cloning vector, which introduced hyperactive E54K and L372P mutation into wildtype Tn5, was acquired from Addgene. The pTXB1 Tn5 and its mutant were expressed and purified mainly according to the protocol published by Picelli S et al. [1] To construct D188E mutation into Tn5, pTXB1 vector was firstly amplified into two parts by two sets of primers. Mutagenesis primers used for the first part (3771-7979) which contained site 188 were 5’-GGCAGCATGATGAGCAACGTGATTGCGGTGTGC**GAA**CG TGAAGCGGATATTCATGC-3’ and 5’-TATCAGCTCACTCAAAGG-3’. Amplified primers for the remaining part were 5’-GTATTACCGCCTTTGAGT-3’ and 5’-CAATCACGTTGCTC ATCA-3’. The purified PCR products were then assembled into intact plasmid using Gibson Assembly Master Mix (NEB, Cat.No. E2611). The newly assembled plasmid was transformed into E.coli Trans5α chemically competent cells (Transgene, CD201-01). After growing overnight on LB medium plate, single colony was picked and shaken in SOC liquid medium for at least 9 hours. Plasmid was extracted by PurePlasmid Mini Kit (CWBIO, Cat.No.CW0500S) and confirmed carrying D188E mutation by Sanger sequencing. Then the plasmid was transformed into E.coli Transetta (DE3) chemically competent cells (Transgene, CD801-01) for further protein expression and purification.

### Cell culture

HEK293T and HeLa cell lines are acquired from ATCC. Both of them were cultured in Dulbecco’s Modified Eagle Medium (Gibco, Cat.No.11965092), supplemented with 10% fetal bovine serum (Gibco, Cat.No.1600044) and 1% penicillin-streptomycin (Gibco, Cat.No.15140122). The cell incubator (Thermo Scientific) was set at temperature of 37°C with 5% CO_2_ injected.

Adherent cells were washed twice by DPBS (Gibco, Cat.No.14190136) and detached by 0.05% Trypsin-EDTA (Gibco, Cat.No.25300062) at 37°C for 4min. Then double volume of culture media was added to terminate trypnization. Cells were collected by centrifugation at 200g for 5min and resuspended for downstream experiment or passage cultivation.

### Nucleic acids extraction and messenger RNA isolation

Genomic DNA was extracted using PureLink Genomic DNA Mini Kit (Invitrogen, Cat.No.K182002), and total RNA was extracted using RNeasy Mini Kit (Qiagen, Cat.No.74104). The resulting total RNA was then reacted with 10ul DNase I (NEB, Cat.No.M0303) to remove remaining DNA thoroughly, and concentrated by RNA Clean & Concentrator-5 kit (Zymo Research, Cat.No.R1015). The quality of extracted DNA and RNA was assessed by the Fragment Analyzer Automated CE System (AATI) and quantification was done by Qubit 2.0 (Invitrogen, Cat.No.Q33230/ Q32852).

We followed standard protocol of NEBNext Poly(A) mRNA Magnetic Isolation Module (NEB, Cat.No.E7490) to isolate messenger RNA from the purified total RNA and stored them at −80°C.

### Single cell preparation

We used pipette tips to form some drops made up of PBS (containing 1% BSA) (Thermo Scientific, Cat.No.37525) on a clean petri dish. The cell resuspension was pipetted up and down gently to disperse into single cells and we took ∼5μl of them diluted in one of the drops. The mouth pipette with 50μm inside diameter was then used to pick one cell in the drop and release it in another clean drop. The picked cell was passed by at least three clean drops in order to wash away any debris and confirm that only one cell was in the last drop. We then aspirated the cell with as little buffer as possible and blew it into 4μl lysis buffer [4 units of Recombinant RNase Inhibitor (Takara, Cat.No.2313), 2.5μM poly(T)_30_VN primer (Sangon), 2.5mM dNTP (NEB, Cat.No.N0447) and 0.48% Triton X-100 (Sigma, Cat.No.T9284)]. Successful transfer was confirmed by blowing mouth pipette again in a clean drop and no cell was to be seen in visual field. Reaction was carried out at 72°C for 3min after violent vortex.

### mRNA/DNA hybrid formation

Total RNA was reverse transcribed into mRNA/DNA hybrid mainly referring to Smart-seq2 protocol [2], but with several modifications: 1) The ISPCR part in Oligo-dT primer was removed; 2) TSO was omitted, but in TSO-RT SHERRY, it should be kept; 3) The reaction was performed at 42°C for 1.5h without cycling. If input was purified RNA, Triton X-100 was omitted. When inputting more than 10ng total RNA, we would slightly upregulate amount of dNTP, poly(T)_30_VN primer and Superscript II (Invitrogen, Cat.No.18064014).

### Tn5 transposome in vitro assembly and tagmentation

Functional mosaic-end (ME) oligonucleotides (5’-CTGTCTCTTATACACATCT-3’, 100μM) was separately annealed with equal amounts of Adaptor A (5’-TCGTCGGCAGCGTCAG ATGTGTATAAGAGACAG-3’, 100μM) and Adaptor B (5’-GTCTCGTGGGCTCGGAGATG TGTATAAGAGACAG-3’, 100μM). The concentration of purified Tn5 was quantified by Qubit Protein Assay Kit (Invitrogen, Cat.No.Q33212) and took around 100μg for transposome assembly. We mixed the Tn5 transposase with annealed ME-Adaptor A or B (20μM) in 45% glycerol (Sigma, Cat.No.G5516) thoroughly and incubated the mixture at 30°C for one hour. These two resulting transposomes (assembled with ME-Adaptor A/B) were then mixed together, ready for tagmentation or stored at −20°C. Specifically, to assemble rCrArG Tn5 in **Fig.S4**, ribonucleotide modifications were made on the three terminal bases at 3’-end of Adaptor A/B. And for rG Tn5, the last base at 3’-end of adaptors was modified.

The dsDNA tagmentation was performed in 1xTD buffer [10mM Tris-Cl (pH 7.6, ROCKLAND, Cat.No.MB-003), 5mM MgCl_2_ (Invitrogen, Cat.No.AM9530G), 10% N,N-Dimethylformamide (Sigma, Cat.No.D4551)]. The reaction was incubated at 55°C for 30min.

As for RNA/DNA hybrid, tagmentation was performed in buffer containing 10mM Tris-Cl (pH 7.6), 5mM MgCl_2_, 10% N,N-Dimethylformamide, 9% PEG8000 (VWR Life Science, Cat.No.97061), 0.85mM ATP (NEB, Cat.No.P0756). In SHERRY library preparation, we used 0.05 μl, 0.006 μl and 0.003 μl Tn5 transposome for input of 200 ng, 10 ng and 100 pg total RNA, respectively. scSHERRY also used 0.003 μl Tn5 transposome.

The transposome could be diluted in 1xTn5 dialysis buffer [50mM Hepes (pH 7.2, Leagene, Cat.No.CC064), 0.1M NaCl (Invitrogen, Cat.No. AM9759), 0.1 mM EDTA (Invitrogen, Cat.No.AM9260G), 1 mM DTT, 0.1% Triton X-100, 10% glycerol].The reaction was incubated at 55°C for 30min.

Commercial Tn5 transposomes were available in Nextera XT DNA Library Prep Kit (Illumina, Cat.No.FC-131-1024) and TruePrep DNA Library Prep Kit V2 for Illumina (Vazyme, Cat.No.TD501).

### SHERRY library preparation and sequencing

To construct scSHERRY library, the single cell tagmentation product was mixed well with 4 units of Bst 3.0 DNA Polymerase (NEB, Cat.No.M0374) and indexed common primers (Vazyme, Cat.No.TD202) in 1 x Q5 High-Fidelity Master Mix (NEB, Cat.No.M0492). Then index PCR was performed as follow: 72°C 15min, 98°C 30s, 10 cycles of [98°C 20s, 60°C 20s, 72°C 2min], 72°C 5min. The PCR product was purified with 0.85:1 ratio by VAHTS DNA Clean Beads (Vazyme, Cat.No.N411) and eluted in 30μl nuclease-free water (Invitrogen, Cat.No.AM9937) for another 18 cycles of PCR. When performing high-throughput experiment, each sample could be amplified by 15 cycles, then merged for beads purification and library quality check.

As for purified 10ng or 200ng total RNA input, the tagmentation product was firstly gap-filled with 100 units of Superscript II and 1 x Q5 High-Fidelity Master Mix at 42°C for 15min, then Superscript II was inactivated at 70°C for 15min. When inputting 100pg total RNA, the extension enzyme was replaced with 4 units of Bst 2.0 Warmstart DNA Polymerase (NEB, Cat.No.M0538). Correspondingly, the reaction temperature was upregulated to 72°C and inactivation was performed at 80°C for 20min. After that, indexed common primers were added to perform PCR. We performed 12, 15 and 25 cycles of PCR for input of 200 ng, 10 ng and 100 pg total RNA, respectively.

The resulting library was purified with 1:1 ratio by VAHTS DNA Clean Beads. Quantification was done by Qubit 2.0 and quality check was done by Fragment Analyzer Automated CE System. The sequencing platform we used was Illumina NextSeq 500 or HiSeq 4000.

### NEBNext and SmartSeq2 library preparation

NEBNext RNA-Seq library preparation starting from 10ng and 200ng total RNA was performed using NEBNext Ultra II RNA Library Prep Kit for Illumina (NEB, Cat.No.E7770). Single cell SmartSeq2 library was constructed as previously reported [2].

### Ligation Test and Strand Test

In Ligation Tests, tagmentation products from 200ng HEK293T total RNA were purified by DNA Clean & Concentrator-5 (Zymo Research, Cat.No.D4013) and eluted with 20μl nuclease-free water. After gap-filling with Superscript II, Ligation Test 1 was processed directly to index PCR while product in Ligation Test 2 was digested by 12.5 units of RNase H (NEB, Cat.No.M0297) at 37°C for 20min before index PCR.

In Strand Tests, we used dUTP (Thermo Scientific, Cat.No.R0133), dATP, dCTP, dGTP (NEB, Cat.No.N0446) mix, each of them at equal concentration, to incorporate in cDNA during reverse transcription step. 0.15μl Tn5 transposome was used to tagment the resulting hybrid. Fragments were then column-purified and gap-filled by Bst 2.0 Warmstart DNA Polymerase, and column purification was again applied. For Strand Test 1, 3 units of USER enzyme (NEB, Cat.No.M5505) and 40units of recombinant rnase inhibitor was added into elution products and incubated at 37°C for 20min for DNA strand digestion. Indexed common primers added with digestion product in 1 x Q5 High-Fidelity Master Mix were then reacted at 85°C for 30s, followed by 60°C for 2min and temperature went down to 4°C slowly. After that, reverse transcription was performed with 200 units of Superscript II added at 42°C for 30min, then transferred to index PCR program. For Strand Test 2, the USER digestion product was directly performed index PCR with 1 x KAPA HiFi HotStart Uracil+ ReadyMix (Kapa Biosystems, Cat.No.KK2801). Protocol of Strand Test 3 was almost same as Strand Test 2, except replacing USER enzyme with RNase H. For dU-SHERRY, the USER enzyme digestion was omitted compared with Strand Test 2 workflow.

### Docking model

To generate the substrate-transposon DNA-Tn5 structure model, a 32bp dsDNA or DNA/RNA hybrid were generated by 3D-NuS (3-Dimensional Nucleic Acid Structures) web server. Then the substrate dsDNA or DNA/RNA was manually docked to the transposon DNA-Tn5 structure, PDB ID: 1MUS, based on charge and shape complimentary.

### Data analysis

Sequencing adaptors or poly(T/A) positioned at end of paired reads were recognized and removed by Cutadapt v1.15 [3]. The trimmed reads which length was shorter than 20bp were filtered. Remaining reads were down sampled to 2 million (except that library with 200ng total RNA input used for differential gene expression analysis was 10 million) total reads, and aligned with index built from human(hg38) genome and known transcript annotations by Tophat2 v2.1.1 [4]. The mapped reads were then used to calculate FPKM value for each known gene (annotation acquired from UCSC) by Cufflinks v2.2.1 with multi-mapped reads correction. Gene with FPKM more than 1 was considered to be detected. The exonic rate, duplicate rate and insert size of library were all calculated by Picard Tools v2.17.6.

General coverage across known transcripts was plotted by RSeQC v.2.6.4 [5]. For specific transcript, depth of mapped reads overlapped with transcript position was calculated by Samtools v1.3.1.

We used DESeq2 v1.22.2 [6] to perform differential gene expression analysis with raw count-matrix acquired by HTSeq v.0.11.0 [7]. Differentially expressed genes should meet following criteria: 1) FPKM value >1; 2) significant p-value <5×10^−6^; 3) absolute value of log_2_(Fold Change) >1. Counts in correlation plot were normalized mainly according to DESeq2 normalization method [6], which considered library size and library compensation. Correlation efficient R^2^ and slope of linear fitting equations were calculated by least square method. The slope-skewness between two replicates was defined as |k_1_-1|+|k_2_-1|, k_1_ or k_2_ was slope when one of the replicates was conducted as X or Y axis.

GC content distribution of detected genes was plotted by custom Perl script. GC content was binned by 2% and gene number at each bin was normalized by dividing maximum gene number of one bin. And for each gene, only the longest transcript isoform was calculated.

### Data deposition

The sequence reported in this paper has been deposited in the Genome Sequence Archive (accession no. CRA002081).

